# Unraveling and overcoming the ammonia toxicity in methanotrophs for sustainable biomanufacturing and methane removal

**DOI:** 10.1101/2025.01.28.635327

**Authors:** Haili Zhang, Zixi Gao, Xi Xiao, Minggen Cheng, Qiang Fei, Xin Yan

## Abstract

Replacement of nitrate with ammonium at large scale cultivation of methanotrophs can improve the economic feasibility of these bacteria in methane-based biomanufacturing and methane removal. However, ammonia toxicity and N_2_O emission impede this option. The mechanism of ammonia oxidation in methanotrophs remains elusive, limiting the effort to detoxify ammonia via genetic engineering. Using an industrially promising methanotroph as a model, we identified a porin PorA that facilitated ammonium uptake. Inactivation of PorA remarkably relieved ammonia toxicity and reduced N_2_O production. Meanwhile, we demonstrated that *haoA*, *cytL* and *hcp* contributed to ammonia detoxification and *cytL* was involved in the conversion of NH_2_OH to N_2_O. A mutant strain with increased ammonium-utilizing ability and decreased N_2_O emission was constructed. High growth rate and cell biomass were achieved in fed-batch fermentation with this strain using ammonium. These results deepen our understanding of ammonia oxidation in methanotrophs and promote their applications in biomanufacturing and methane removal.

**IMPORTANCE:** *M. buryatense* 5GB1C is becoming a methanotrophic model for both fundamental research and practical applications. Here, the genetic determinants of ammonia toxicity were identified and a porin gene that facilitated ammonium uptake was described for the first time, providing new understanding regarding the mechanism of nitrification in methanotrophs. Moreover, this study dramatically reduced ammonia toxicity and N_2_O production in this strain and enabled the use of ammonium as nitrogen source in fermentation, improving the sustainability of this strain in practical applications.

## INTRODUCTION

Aerobic methanotrophic bacteria are essential players in the global ecosystem by utilizing methane as their primary carbon and energy source under ambient conditions. These bacteria are widely distributed in various natural environments and serve as critical biofilter of methane (1). In addition to their ecological importance, methanotrophs have drawn attention for their potential in industrial applications, particularly in methane-based biomanufacturing and as biocatalysts for methane removal at emission sites (2–5). Valuable products including chemicals, natural products, and biopolymers, biofuels (6–10) have been synthesized by natural and genetic engineered methanotrophs. Meanwhile, significant progress has been made on methanotroph-based methane removal technology (2). Economic and environmental feasibility are crucial factors affecting the commercial success of methanotrophs-based industrial applications (11). Nitrogen source employed in biomanufacturing significantly impacts overall production cost. Since the price of ammonium is usually the half of that of nitrate, replacement of nitrate with ammonium as nitrogen source has been used previously as a strategy to reduce the cost of various fermentation processes (12, 13). However, due to the toxicity of ammonia to methanotrophs, these bacteria are traditionally grown using nitrate as nitrogen source (14–17). Additionally, since N_2_O is generated during ammonia oxidation in methanotrophs (14), N_2_O emission should be diminished when using ammonia as nitrogen source.

Genetic engineering is one potential solution to enhance the ability of methanotrophs to utilize ammonium. Nevertheless, the underlying mechanisms of ammonia toxicity in methanotrophs are not yet fully understood, posing challenges for effective genetic manipulation. Due to their evolutionary relatedness to ammonia-oxidizing bacteria, methanotrophs are usually capable of oxidizing ammonia to generate hydroxylamine (NH_2_OH), nitric oxide (NO), nitrous oxide (N_2_O) and nitrite (NO_2_ ^-^) (18, 19). The negative effects of ammonia on methanotrophs can be largely attributed to the competitive inhibition of methane monooxygenase as well as the toxic properties of intermediates such as NH_2_OH, NO or NO_2_^-^ (17). This highlights a critical area of research that could pave the way for advancements in methanotrophic applications.

The genetic and biochemical background of ammonia oxidation in ammonia-oxidizing bacteria is shown in FIG. 1A. In these microorganisms, ammonia is oxidized to NH_2_OH by ammonia monooxygenase (AMO) (20, 21). Subsequent reactions involve hydroxylamine oxidoreductase (HAO), which converts NH_2_OH to NO (22), while cytochrome P460 (CytL) further transforms NH_2_OH to NO_2_^-^ and N_2_O under aerobic condition(23). Additionally, Kozlowski et al. reported that a membrane-bound nitric oxide reductase (NorB) plays a role in the N_2_O production via hydroxylamine oxidation pathway in *Nitrosomonas europaea* ATCC 19718 (24). In methanotrophs (FIG. 1B), particle methane monooxygenase (pMMO) is evolutionarily related to AMO, and both enzymes can oxidize ammonia to NH_2_OH (25, 26). The homologs of HAO, CytL and NorB have been found in some methanotrophs (27–31). However, their specific roles in ammonia oxidation remain to be validated via genetic or biochemical evidence (27, 28, 30–33).

**FIG 1.**
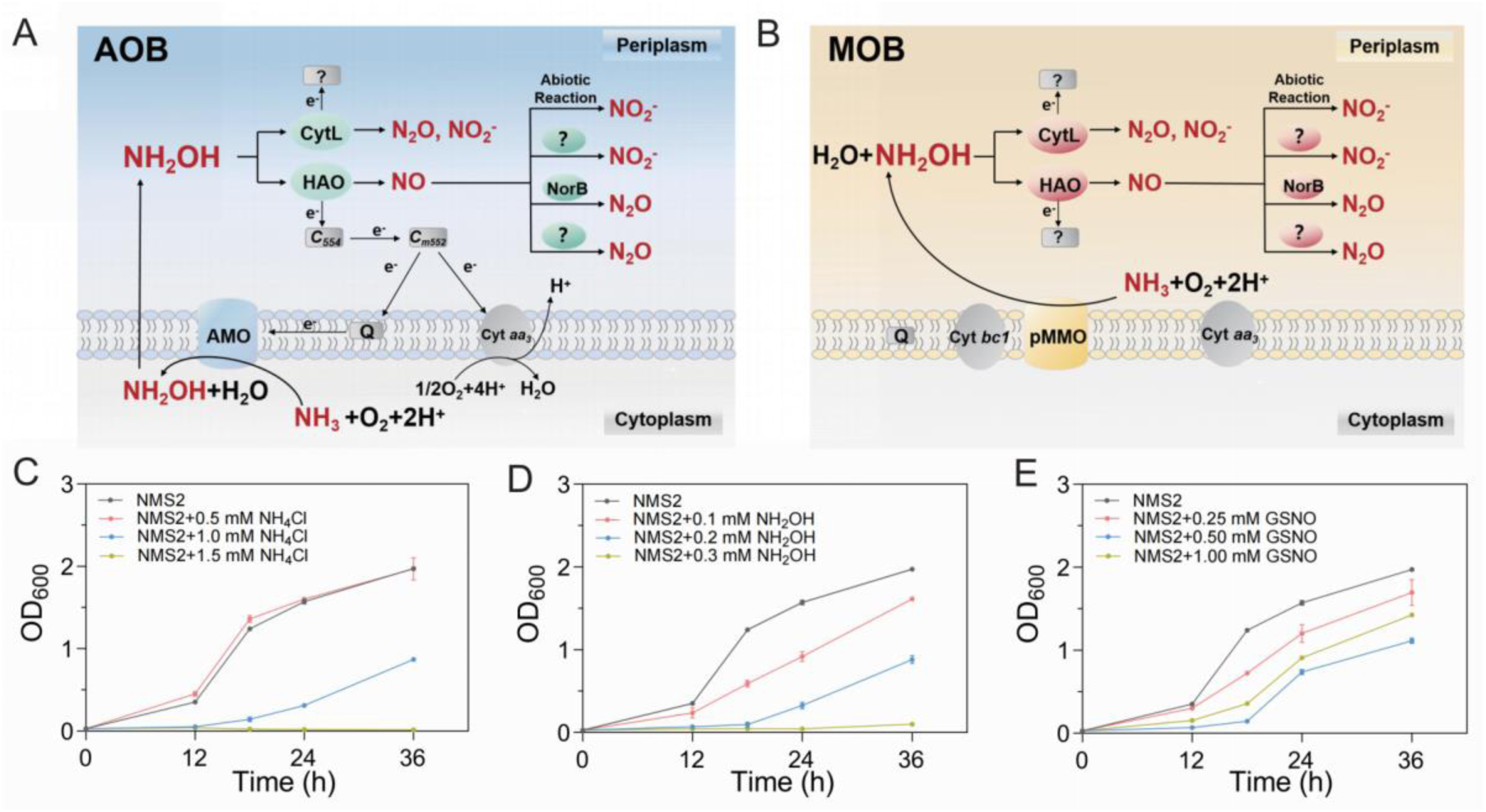
AOB and MOB mediated ammonia oxidation and the sensitivity of *M. buryatense* 5GB1C to ammonium and its oxidized products. (**A)** Proposed metabolic pathway for ammonia oxidation in AOB. (**B)** Proposed metabolic pathway for ammonia oxidation in MOB. (**C), (D), (E)** Growth curves of *M. buryatense* 5GB1C in NMS2 medium supplemented with different concentrations of NH_4_Cl, NH_2_OH and S-nitrosoglutathione, GSNO. GSNO is an amino acid nitric oxide donor that releases nitric oxide (NO) under physiological conditions. All growth test were performed with three biological replicates, and error bars represent SDs (*n*=3).

Among the studied methanotrophs, *M. buryatense* 5GB1C stands out as a promising candidate for both biomanufacturing and methane removal. This gammaproteobacterial methanotroph, isolated from extreme environment of soda lake sediments (34), demonstrates robust growth on natural gas and exhibits heightened resistance to contamination. Efficient genetic tools and increasing fundamental knowledges make *M. buryatense* 5GB1C as an ideal chassis for industrial applications (35–39). Moreover, He et al. demonstrated that *M. buryatense* 5GB1C is a candidate to develop technology for methane removal at emission sites, owing to its outstanding ability to grow and consume low concentration methane (2). However, despite possessing the genes for HAO, CytL and NorB, this strain is highly sensitive to ammonium, and N_2_O is generated in the presence of ammonium. Although *M. buryatense* 5GB1C can use ammonium at lower pH levels (around 7.0), its growth rate significantly declines, with doubling time extending from 3 to 12 hours (14).

In this light, this study aims to: i) identify the genes associated with ammonia toxicity in *M. buryatense* 5GB1C; ii) enhance the ammonium-utilizing ability of this strain through genetic engineering; iii) minimize N_2_O emissions under ammonium condition; iv) develop culture strategy to realize high growth rate and cell biomass with ammonium. Through these efforts, we hope to contribute to the advancement of methanotroph-based practical applications.

## RESULTS

### The Sensitivity of *M. buryatense* 5GB1C to Ammonium and Its Intermediates

Initially, the sensitivity of *M. buryatense* 5GB1C to ammonium and its potential oxidation products was investigated. This strain is usually grown in nitrate mineral salt medium (NMS2) that contains 10.0 mM KNO_3_ as the sole nitrogen source (pH 9.5). Addition of 1.0 mM NH_4_Cl to NMS2 medium dramatically inhibited cell growth, with a lag phase of approximately 16 hours. When 1.5 mM NH_4_Cl was added, no cell growth was observed within 36 hours (FIG. 1C). NH_2_OH also severely impeded cell growth and no cell growth was detected within 36 hours in the presence of 0.1 mM NH_2_OH (FIG. 1D). The sensitivity to NO was evaluated using S-nitrosoglutathione (GSNO), with obvious growth inhibition occurred at a concentration of 0.25 mM (FIG. 1E). In contrast, addition of NaNO_2_ (3.0 mM and 5.0 mM) did not affect cell growth (data not shown).

When the *M. buryatense* 5GB1C was cultured in ammonium mineral salt medium (AMS2), a growth rate slightly faster than that with 0.5 mM KNO_3_ was observed at 0.5 mM NH_4_Cl. However, 1.0 mM NH_4_Cl resulted in an extended lag phase of 24 hours, and 1.5 mM NH_4_Cl caused complete growth arrest within 48 hours (FIG. S1A). Similarly, *M. buryatense* 5GB1C could utilize NH_2_OH as the sole nitrogen source for growth only when it was supplied at low concentration of 0.1 mM (FIG. S1B). In contrast, the strain grew efficiently when NaNO_2_ was provided as the nitrogen source (FIG. S1C). These results indicated that the toxicity of ammonium, NH_2_OH and NO could impair the ammonium-utilizing ability of *M. buryatense* 5GB1C.

### Screening of a Transposon Mutagenesis Library for Mutants with Enhanced Ammonium-Utilizing Ability

To explore new genes associated with ammonia toxicity, we established an efficient transposon mutagenesis method for *M. buryatense* 5GB1C. As shown in FIG. 2A, the transposon library was screened on the plates of NMS2 and NMS2+NH_4_Cl plates, respectively. The mutant with elevated growth rate on NH_4_Cl-containing plate was selected. One such mutant, named 5G-M1, was obtained, which exhibited outstanding capability of growing on NH_4_Cl-containing NMS2 plate (FIG. 2B). Arbitrary PCR revealed that gene EQU24_19165 was destroyed by transposon in mutant 5G-M1 (FIG. 2C). The results of gene knockout and genetic complementation confirmed that inactivation of EQU24_19165 enhanced the ammonium-utilizing ability of *M. buryatense* 5GB1C on agar plates (FIG. 2D). Notably, mutant Δ*19165* and wild-type strain shared similar growth curves when KNO_3_ was used as the sole nitrogen source (FIG. 2E). After transferred from KNO_3_ condition to NH_4_Cl condition, mutant Δ*19165* grew well in the liquid medium with 1.0 mM or 2.0 mM NH_4_Cl, but exhibited an extended lag phase of approximately 40 hours with 3.0 mM NH_4_Cl (FIG. 2F). These results showed that gene EQU24_19165 remarkably affected the ammonium-utilizing ability of *M. buryatense* 5GB1C, but had little effect on its nitrate-utilizing ability.

**FIG 2.**
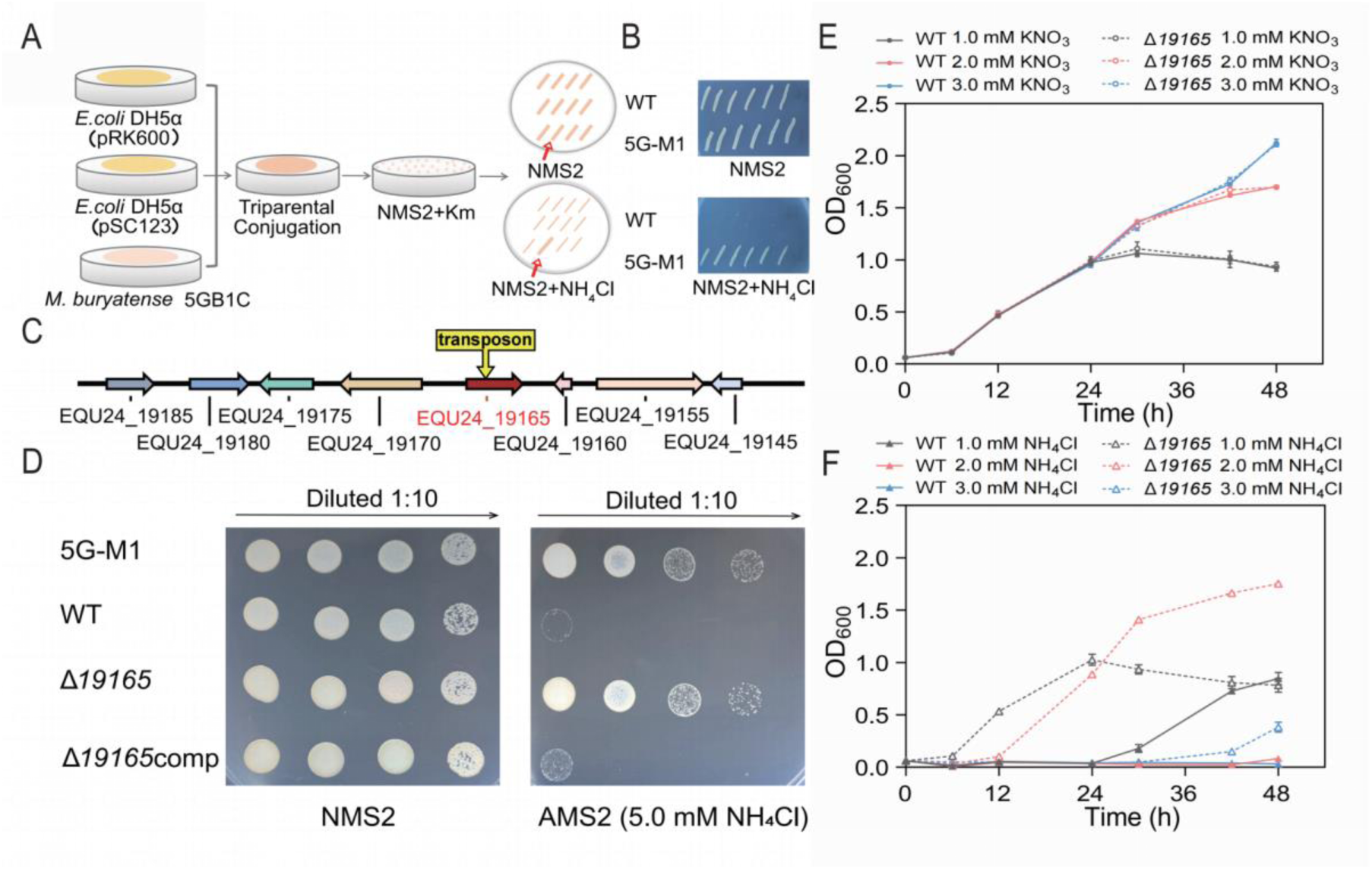
Screening of transposon mutagenesis library for the genes involved in ammonia toxicity. (**A)** Construction and screening of transposon mutagenesis library. (**B)** Mutant strain 5G-M1 with enhanced ammonium-utilizing ability. (**C)** Transposon insertion site in mutant strain 5G-M1. (**D)** Verify the role of gene EQU24_19165 via gene knockout and genetic complementation. Each culture was serially diluted and equal amounts of cells were spotted onto NMS2 and AMS2 plates. Δ*19165*, EQU24_19165 deletion mutant; Δ*19165*comp, genetic complementation strain of Δ*19165*. (**E)** Growth curves of mutant strain Δ*19165* in liquid media with NH_4_Cl or KNO_3_ as nitrogen source. Error bars represent SDs (*n*=3).

### EQU24_19165 Encodes a Putative Porin that Affected the Absorption of Total Ammonia Nitrogen (TAN)

The product of gene EQU24_19165 was predicted to be a porin, which consists of 402 amino acids, with a 26-aas signal peptide at its N-terminal. The result of AlphaFold 3 prediction showed that this protein displayed a typical porin structure, with 16 β-strands that were arranged into a β-barrel (FIG. 3A). Gene EQU24_19165 was named *porA* and mutant Δ*19165* is equal to Δ*porA*. Porins are typically located in the outer membrane of Gram-negative bacteria and serve as channels for small molecules such as ions and antibiotics (40). Antibiotic-specific porins enhance antibiotic uptake, thereby increasing susceptibility (41). Considering that *porA* affected the utilization of ammonium but not nitrate, we tested whether *porA* is involved in TAN uptake. Notably, TAN includes both ammonia (NH_3_) and ammonium (NH_4_^+^). The cells grown in liquid NMS2 were harvested, washed and then suspended in the medium containing 0.3 mM NH_4_Cl. The TAN in cell free supernatant was measured with time. As shown in FIG. 3B, the TAN concentration in Δ*porA* treatment was significantly higher than that in the wild-type strain at the same time points, indicating that inactivation of *porA* decreased TAN uptake efficiency. Considering that reduced TAN uptake can result in less production of NH_2_OH and NO_2_^-^, the concentrations of these two compounds in cell free supernatant were analyzed in the presence of 1.0 and 2.0 mM NH_4_Cl. As expected, knockout of *porA* remarkably decreased the production of both NH_2_OH (FIG. 3D) and NO_2_^-^ (FIG. 3D).

**FIG 3.**
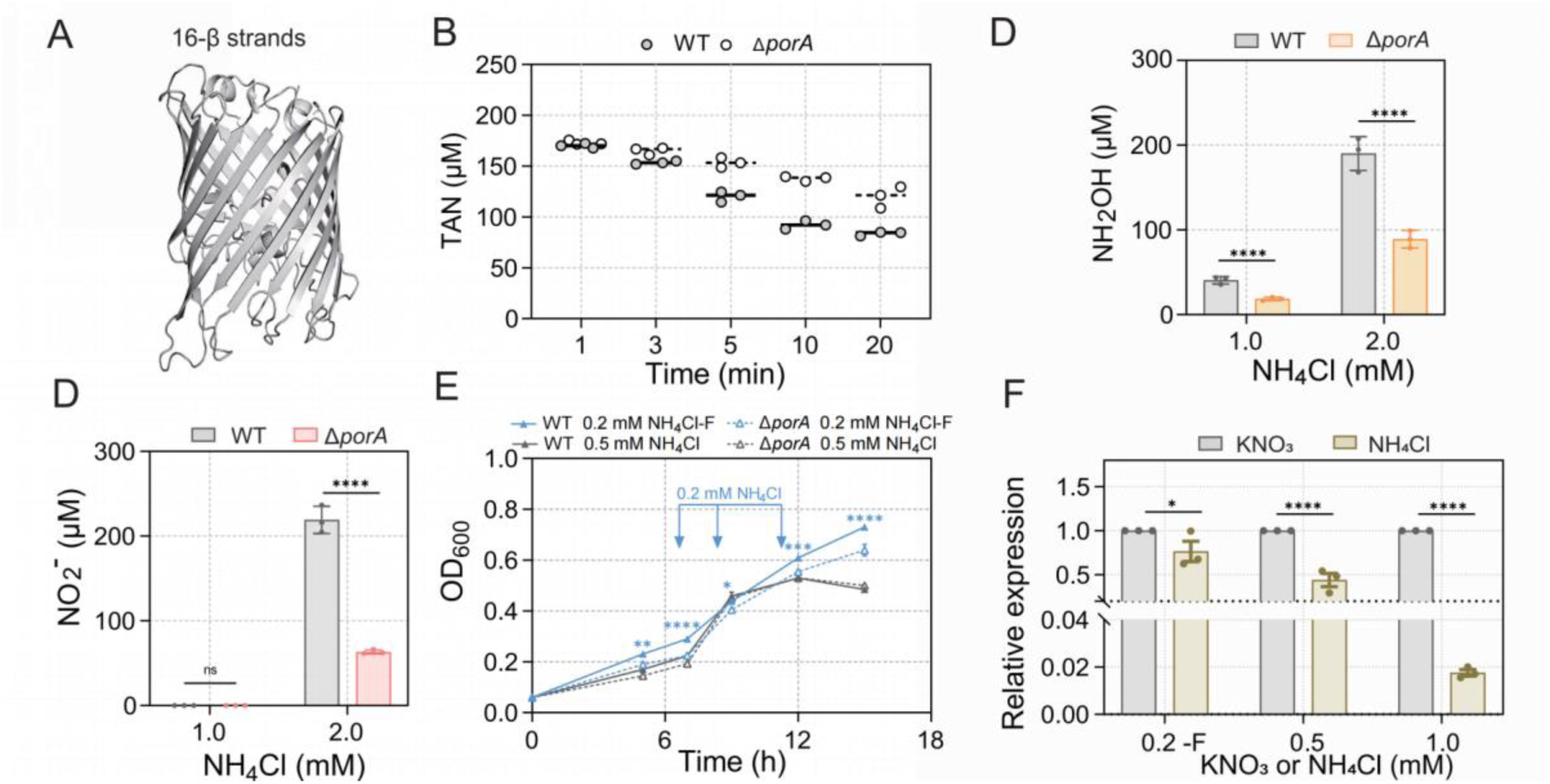
*porA* was involved in the uptake total ammonia nitrogen (TAN). (**A)** Side view of a PorA monomer predicted by AlphaFold3. (**B)** TAN adsorption capacity of wild-type strain and mutant strain Δ*porA* (also known as EQU24_19165) (**C), (D)** The production of NH_2_OH and NO_2_^-^ in wild-type strain and mutant strain Δ*porA* using NH_4_Cl at nitrogen source. **(E)** Growth curves of wild-type strain and mutant strain Δ*porA* at lower NH_4_Cl concentrations. “0.2 F” represents that additional 0.2 mM NH_4_Cl was supplemented at the time points indicated with blue arrows. (**F)** Transcriptional analysis of *porA* via RT-qPCR. The cells were cultivated to the late exponential phase with NH_4_Cl or KNO_3_ as nitrogen sources. *ns,* no significant difference; \**P* < 0.05; \*\**P* < 0.01; \*\*\**P* < 0.001; \*\*\*\**P* < 0.0001. Significance test was determined by student’s *t*-test or two-way ANOVA. Error bars represent SDs (*n*=3).

It is unreasonable for a bacterium to express a porin that hampers its TAN utilization in the natural environment. Therefore, we tested whether *porA* is beneficial to *M. buryatense* 5GB1C at lower TAN concentrations, which are close to the case in the natural environment. Compared to mutant Δ*porA*, parent strain exhibited significantly higher growth rate at 0.2 mM NH_4_Cl, and slightly higher growth rate at 0.5 mM NH_4_Cl (FIG. 3E), which was opposite to the results at 1.0 and 2.0 mM NH_4_Cl (FIG. 2F). To determine the transcriptional response of *porA* to NH_4_Cl via RT-qPCR, cells was transferred from KNO_3_ condition to NH_4_Cl condition and grown to the middle of log phase. Compared to KNO_3_ condition, the transcription of *porA* was down-regulated 1.4, 2.4 and 57.6 times at the 0.2, 0.5 and 1.0 mM NH_4_Cl, respectively (FIG. 3F), which were consistent with the roles of *porA* at these concentrations (FIG. 3E; FIG. 2F). These data showed that expression of *porA* facilitated TAN uptake, which was advantageous at lower ammonium concentrations but disadvantageous under higher ammonium concentrations.

### Identification of the Genes involved in the Detoxication of NH_2_OH and NO

To further improve the ammonium-utilizing ability of mutant strain Δ*porA via* genetic engineering, we identified the genes involved in the detoxication of NH_2_OH and NO. The candidate genes were predicted from the genome of *M. buryatense* 5GB1C. Apart from the genes for HAO (*haoA*), CytL (*cytL*) and NorB (*norB*) (FIG. 4A), *hcp* that encodes hybrid cluster protein (Hcp) was also selected. Hcp was previously reported to convert NH_2_OH to NH_3_ (42), but recent study suggested that its possible physiological role in *E.coli* is converting NO to N_2_O under anaerobic condition (43, 44). RT-qPCR analysis was performed using the cells transferred from KNO_3_ medium to NH_4_Cl medium, which revealed that the transcription of these three genes was significantly upregulated at early log phase but the upregulated amplitude was decreased in the late log phase (FIG. 4B). Deletion the gene of *haoA*, *cytL*, *norB* or *hcp* had little influence on cell growth under KNO_3_ condition. By contrast, all the mutants of Δ*haoA*, Δ*cytL* and Δ*hcp* became more sensitive than the parent strain under NH_4_Cl condition (FIG. 4C-E), suggesting their involvement in ammonia detoxification. In contrast, deletion of *norB* did not affect cell growth under NH_4_Cl conditions (data not shown).

**FIG 4.**
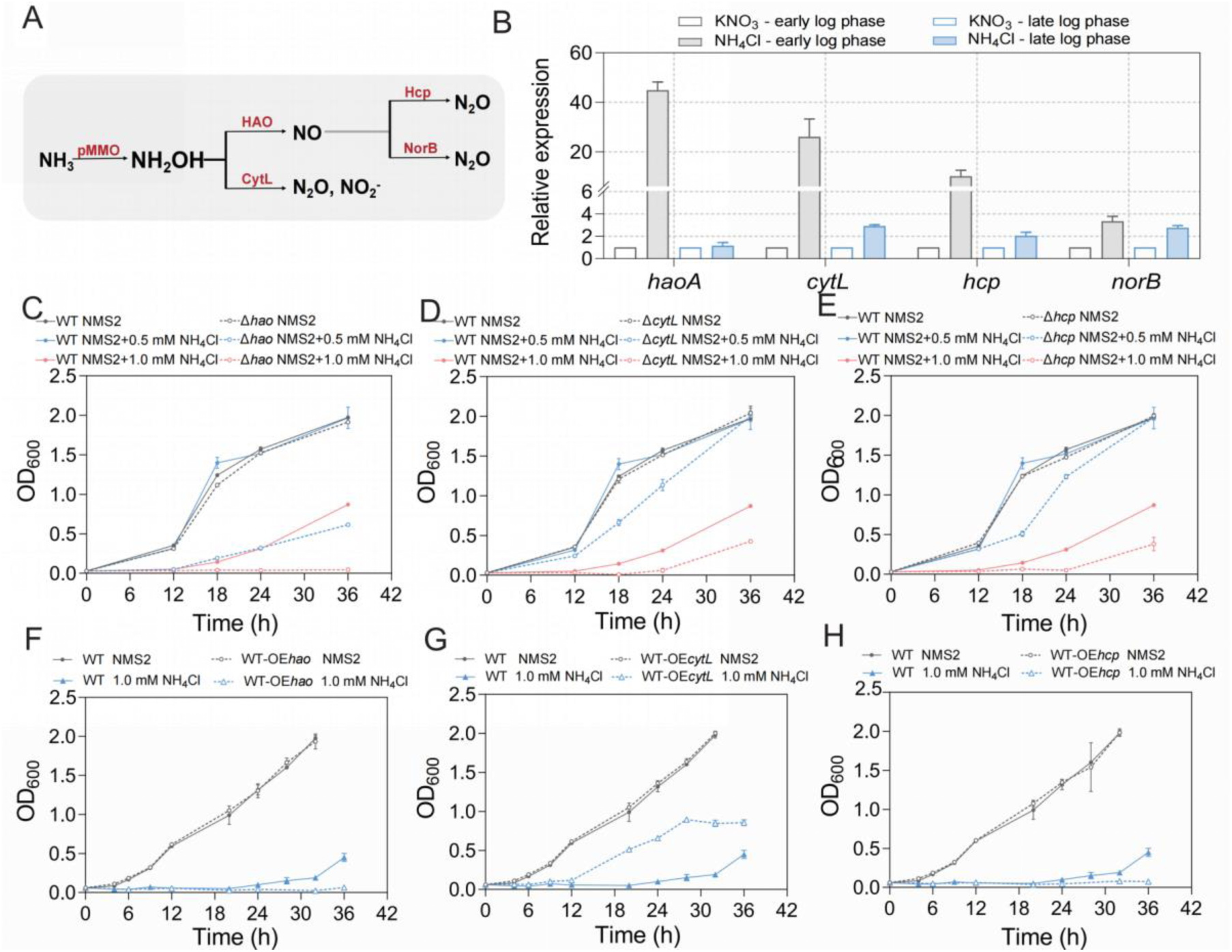
Identification of the genes involved in ammonia detoxication. (**A)** Prediction of the enzymes involved in ammonia detoxication for the genome of *M. buryatense* 5GB1C. (**B)** Transcriptional analysis of *haoA*, *cytL*, *hcp*, and *norB* in wild-type strain. The cells grown under KNO_3_ conditions was transferred to media supplemented with NH_4_Cl or KNO_3_ as nitrogen source. Samples were collected at early and late exponential phases. The mRNA levels under nitrate condition were used as the control. (**C), (D), (E)** The Growth curves of the mutant strains Δ*haoA*, Δ*cytL*, and Δ*hcp* in the NMS2, *cytL* or *hcp*. Error bars represent SDs (*n*=3).

To improve ammonia detoxication, *haoA*, *cytL* and *hcp* were overexpressed individually in both wild-type strain and mutant strain Δ*porA* using the constitutive promoter *P_tac_* (36). Under KNO_3_ condition, overexpression each of these three genes did not affect cell growth. Under NH_4_Cl condition, only *cytL* overexpression was beneficial, while overexpression of either *haoA* or *hcp* resulted in severe growth inhibition (FIG. 4F-H; FIG. S2A-C). As described in the following results, CytL could convert NH_2_OH to N_2_O, so overexpression of *cytL* is not an ideal option to enhance ammonia detoxication.

### CytL was Involved in the Production of N_2_O in the Presence of Ammonium

Next, we identified the roles of genes *cytL*, *hcp* and *norB* in N_2_O generation under ammonium condition. Deletion of *cytL* significantly reduced the production of N_2_O under ammonium conditions (FIG. 5A), while knockout of either *hcp* or *norB* had little impact on N_2_O production (data not shown). To confirm its function, CytL-6His was expressed in *E. coli* and purified (FIG. 5B). The purified CytL-6His could converted NH_2_OH to N_2_O and NO_2_^-^ under aerobic condition (FIG. 5C, D). These results suggested that *cytL* was involved in the production of N_2_O in the presence of ammonium.

**FIG 5.**
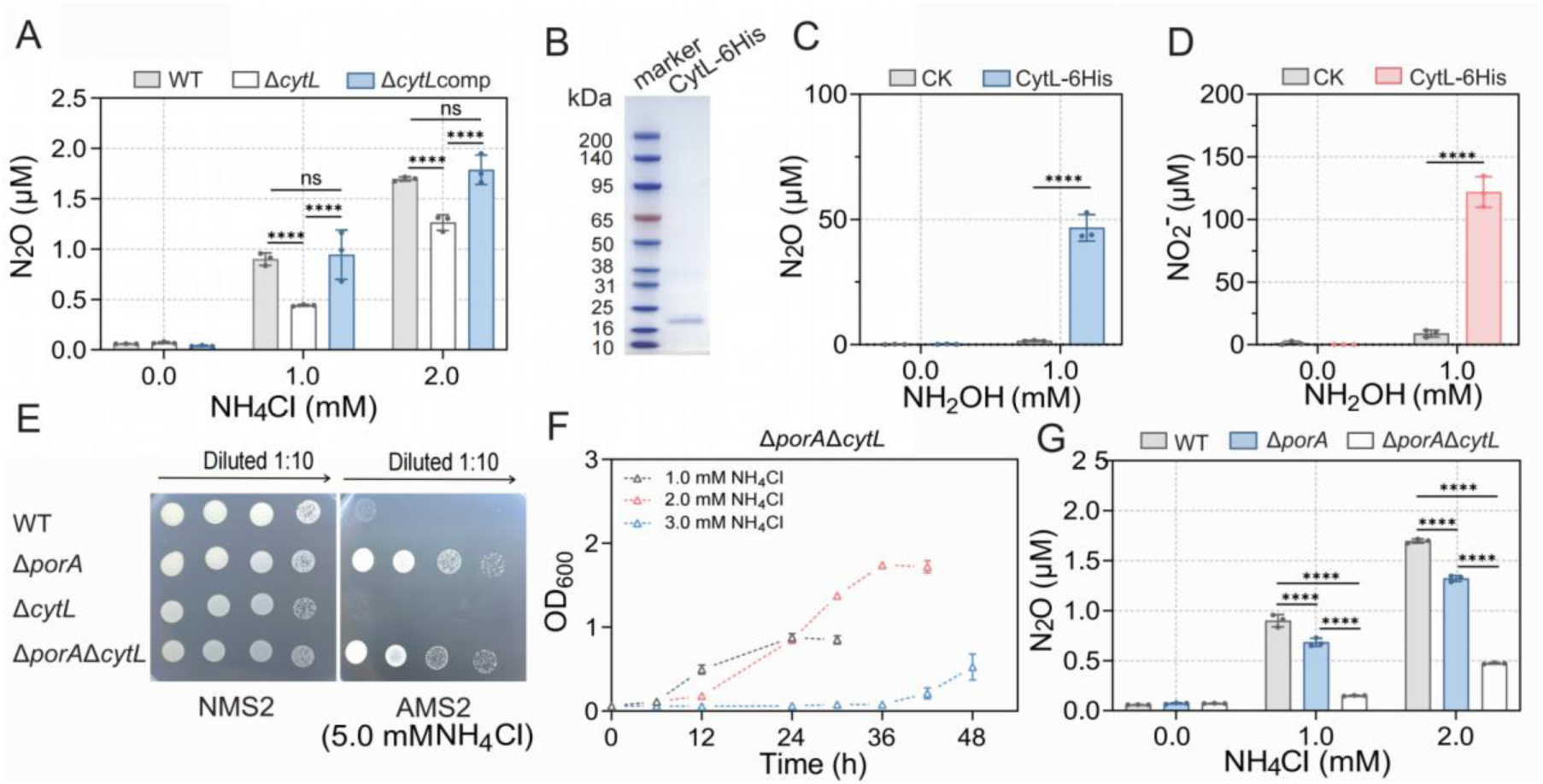
Deletion of *porA* or *cytL* reduced the production of N_2_O. (**A)** Genetic evidence for the involvement of *cytL* in N_2_O production under ammonium condition. (**B)** The purified recombinant CytL-6His expressed in *E. coli* BL21(DE3) Δ*degP*. (**C), (D)** CytL-6His concerted NH_2_OH to produce N_2_O and NO_2_^-^ under aerobic conditions. CK represents the Tris-HCl buffer without CytL-6His. (**E)** Growth of mutant strain Δ*cytL* and Δ*porA*Δ*cytL* on the plates with nitrate or ammonium as nitrogen source. (**F)** Growth curves of mutant strain Δ*porA*Δ*cytL* in the medium using NH_4_Cl as the sole nitrogen source. (**G)** N_2_O production in wild-type strain and mutant strain Δ*cytL* and Δ*porA*Δ*cytL* under ammonium conditions. \*\*\*\**P* < 0.0001. Significance test was determined by student’s t-test or two-way ANOVA. Error bars represent SDs (*n*=3).

### The mutant strain Δ*porA*Δ*cytL* displayed enhanced ammonium-utilizing ability and reduced N_2_O production

The double mutants of *porA* and *cytL* was constructed via unmarked knockout. Although deletion of *cytL* increased the ammonium sensitivity of wild-type strain (FIG. 5E), it did not affect the ammonium-utilizing ability of mutant strain Δ*porA* at the tested NH_4_Cl concentrations (FIG. 5E, F). What is noteworthy is that knockout of *porA* could significantly reduce the production of N_2_O in wild-type strain. Deletion of *cytL* further decreased N_2_O generation under Δ*porA* background. The N_2_O production in double mutant strain Δ*porA*Δ*cytL* was less than 30% of that in wild-type strain at 1.0 and 2.0 mM NH_4_Cl (FIG. 5G).

### Cultivation Conditions Optimization for Mutant Strain Δ*porA*Δ*cytL* using NH_4_Cl as nitrogen source

To tested whether pre-cultivation in ammonium could further enhance the ammonium-utilizing ability of *M. buryatense* 5GB1C and its mutant strains, these strains were first grown to log phase with 1.0 mM NH_4_Cl as the sole nitrogen source and then transferred the medium with different concentrations of NH_4_Cl. Compared to the seed from KNO_3_ medium (FIG. 2F and 5F), pre-cultivation in NH_4_Cl medium significantly enhanced the ammonium-utilizing ability of all tested strains (FIG. 6A-C). Mutant strain *ΔporAΔcytL* could grow well at 3.0 mM NH_4_Cl after pre-cultivation.

**FIG 6.**
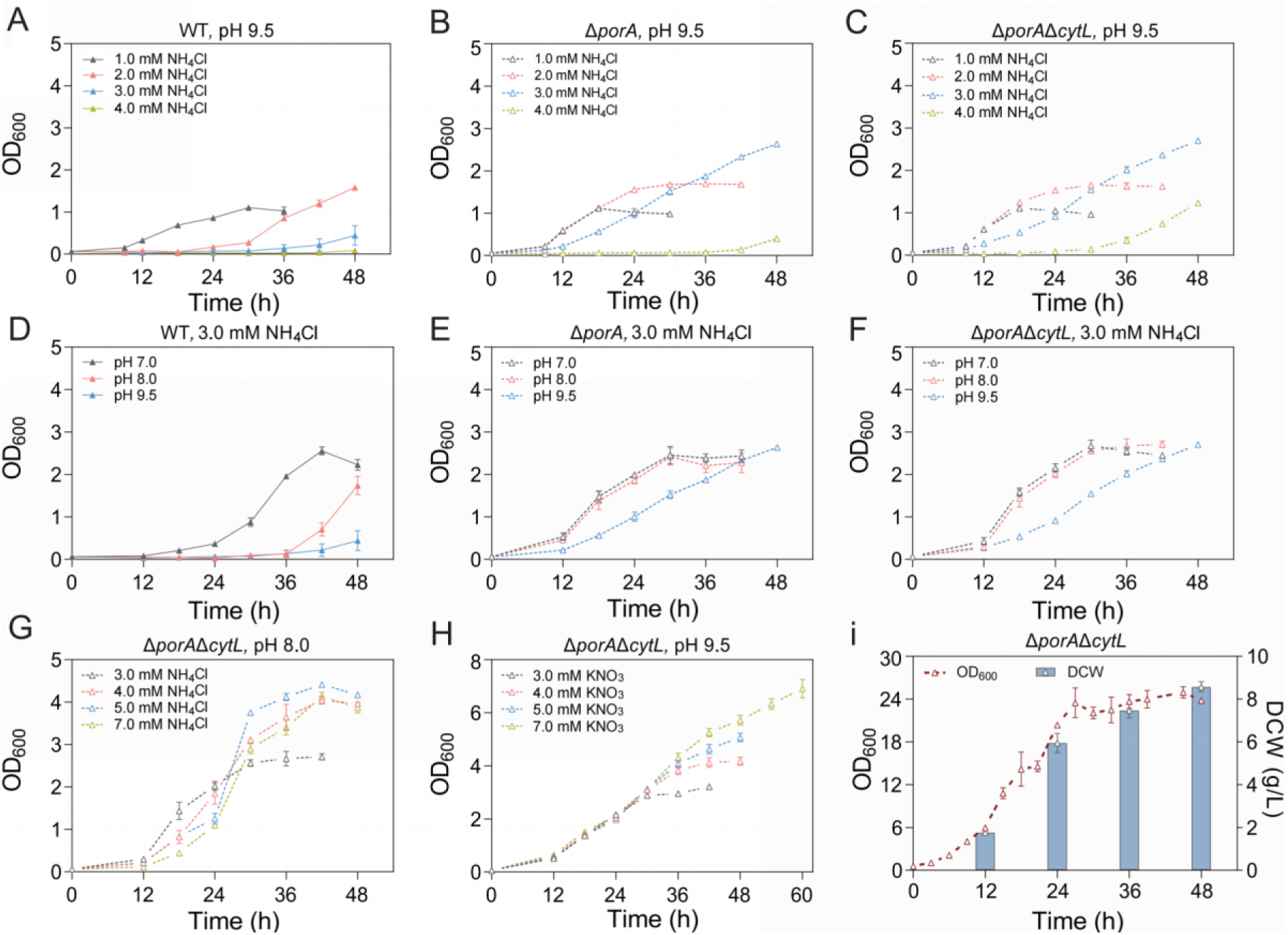
Cultivation conditions optimization for mutant strain Δ*porA*Δ*cytL* using NH_4_Cl as nitrogen source. **(A), (B), (C)** Pre-cultivation in NH_4_Cl enhanced the growth capability of wild-type strain a, Δ*porA* and Δ*porA*Δ*cytL* under ammonium conditions. (**D), (E), (F)** The influence of initial pH on the growth of wild-type strain, Δ*porA* and Δ*porA*Δ*cytL* under ammonium conditions. (**G)** Growth of mutant strain Δ*porA*Δ*cytL* with different concentrations of NH_4_Cl at pH 8.0. (**H)** Growth of mutant strain Δ*porA*Δ*cytL* with KNO_3_ as the nitrogen source at pH 9.5. (**I)** Growth of mutant strain Δ*porA*Δ*cytL* in 3-L fermenters. The seed was pre-cultivated in NH_4_Cl medium. The initial pH value was set at 8.0 and the concentration of TAN was kept at about 3.0 mM. Cell density (OD_600_) and dry cell weight (DCW) are presented as the means ± SD of two 3-L fermenters.

The percent of TAN present as un-ionized ammonia is so dependent upon pH. Previous studies have shown that *M. buryatense* 5GB1C can utilize ammonium as a nitrogen source at a lower pH of 7.0, although with a slower growth rate (14). Therefore, we inoculated NH_4_Cl-induced seed into 3.0 mM NH_4_Cl medium with initial pH values of 7.0, 8.0 and 9.5. Consistent with prior findings (14), lower pH values indeed benefited the growth of all tested strains in NH_4_Cl medium (FIG. 6D-F). Furthermore, deletion of *porA* also dramatically enhanced the ammonium-utilizing ability of *M. buryatense* 5GB1C under pH values of 7.0 and 8.0. The optimal initial pH values for mutants **Δ***porA* and Δ*porA*Δ*cytL* were 7.0 and 8.0, with maximum specific growth rates of 0.220/h and 0.263/h, respectively. We further cultivated mutant strain Δ*porA*Δ*cytL* with increased NH_4_Cl concentration at pH 8.0. The result showed that mutant strain Δ*porA*Δ*cytL* was able to grow under NH_4_Cl concentrations of 3.0, 4.0, 5.0, 7.0 mM (FIG. 6G). It’s worth noting that the growth rate and maximum biomass of mutant Δ*porA*Δ*cytL* under 4.0 mM NH_4_Cl and pH 8.0 were comparable to these in NMS2 medium containing 4.0 mM KNO_3_ (FIG. 6G, H). However, compared to KNO_3_, the utilization efficiency of NH_4_Cl was significantly decreased at the concentrations of 5.0 and 7.0 mM. The maximum OD_600_ at 7.0 mM NH_4_Cl was even lower than that at 4.0 mM NH_4_Cl. These results indicated that the ammonium-utilizing ability of *M. buryatense* 5GB1C could be remarkably improved via the combination of pre-cultivation in ammonium, reduction of initial pH and inactivation of *porA*.

### Replacement of nitrate with ammonium in fed-batch fermentation

Based on the results just above, cultivation of mutant strain Δ*porA*Δ*cytL* was performed in 3-L fermenters using ammonium as the sole nitrogen source. The concentration of NH_4_Cl was maintained at roughly 3.0 mM and the initial pH was set at approximately 8.0. After inoculation with the seed pre-cultivated in ammonium, the cells rapidly entered into the exponential growth phase (FIG. 6I), which resulted in a maximum specific growth rate of 0.225/h (during 3 to 9 h). When stopped the feeding of NH_4_Cl at 24 h, cell growth was slowed down at 27 h and then entered the stagnant phase. In the exponential growth period (from 3 to 24 h), the average specific growth rate was 0.141/h. During the whole fermentation process, approximately 2.14 g/L of NH_4_Cl was utilized and the maximum dry cell weight (DCW) was about 8.57 g/L at 48 h, giving a cell mass productivity and yield of 0.18 g/L/h (16-fold of that in flask culture) (14) and 15.30 g/g N respectively. The total methane utilization was calculated to be 25.01 g (FIG. S3C; Table S3), and thus the cell mass yield on methane was 0.69 g/g, comparable to the continuous cultivation of wild-type strain *M. buryatense* 5GB1 on nitrate as nitrogen source (0.68-0.82 g/g) (7).

## DISCUSSION

Methanotrophs have traditionally relied on nitrate as their nitrogen source due to ammonia toxicity during cultivation. Substitution of ammonium for nitrate could significantly enhance the economic viability of methanotroph-based applications. Furthermore, in formulating emission or cost reduction strategies, it is imperative to concurrently consider the mitigation of CH_4_ and N_2_O emissions (45). In this study, using an industrially promising methanotroph *M. buryatense* 5GB1C as a chassis, we discovered that deletion of the porin gene, *porA*, remarkably improved the ability of this strain to utilize ammonium as nitrogen source. Additionally, the study identified the roles of specific genes encoding for HAO, CytL and Hcp in the detoxification of NH_2_OH. Notably, *cytL* was implicated in the production of N_2_O in the presence of ammonium. The double mutants of *porA* and *cytL* exhibited increased ammonium-utilizing ability and decreased N_2_O production. By pre-cultivating the mutant strain Δ*porA*Δ*cytL* in ammonium, adjusting the initial pH, and systematically feeding ammonium during cultures, we successfully replaced nitrate with ammonium achieving a considerable growth rate. These findings not only deepen our understanding about nitrification in methanotrophs, but also offer pathways for cost reduction and N_2_O mitigation in commercial-scale methanotrophic cultivation.

### The Role of *porA* in Ammonium Utilization

The cell envelope of Gram-negative bacteria is constituted by an outer membrane, the periplasmic space and an inner membrane. Porin proteins on the outer membrane are composed of 8-24 β-strands arranged into β-barrels, which mediate the passive or active uptake of small molecules (40). To the best of our knowledge, no specific porin for ammonium or ammonia has been characterized. Our findings indicate that deletion of *porA* did not hinder cell growth when using nitrate as nitrogen source but dramatically strengthened the ammonium-utilizing ability of *M. buryatense* 5GB1C at the concentration of 1.0 mM or higher (FIG. 2F). Interestingly, compared to wild-type strain, mutant strain Δ*porA* exhibited diminished uptake of TAN (FIG. 3E) and produced lower amounts of NH_2_OH, NO_2_^-^, and N_2_O (FIG. 3C, D; FIG. 5G), suggesting that PorA facilitates the uptake of TAN. TAN consists of a mixture of NH_3_ and NH_4_^+^, with their ratios varying according to pH levels (46). At pH values of 7.0, 8.0, and 9.5, the percent NH_3_ in aqueous ammonia solutions (30°C) were 0.799%, 7.46%, and 71.8%, respectively (47). Deletion of *porA* remarkably enhanced the ammonium-utilizing ability of *M. buryatense* 5GB1C at pH values of 7.0, 8.0, and 9.5 (FIG. 6D-F). At neutral pH (7.0), where approximately 99.2% of TAN is present as NH_4_^+^, it is likely that PorA can promote the passage of NH_4_^+^. However, we cannot exclude the possibility of its involvement in NH_3_ transport.

Both TAN and PorA seem to be double-edged swords to *M. buryatense* 5GB1C. At lower concentration (0.2 mM), TAN posed little adverse effect on cell growth and was therefore a beneficial nitrogen source. Under this concentration, expression of PorA promoted TAN uptake and accelerated cell growth (FIG. 3E), reflecting potential natural environmental interactions. Conversely, at higher concentration of TAN (≥1.0 mM), the existence of PorA resulted in accelerated TAN uptake and more toxic byproducts that the cell failed to cope with. Consequently, the inactivation of PorA appears to be a mechanism to slow down TAN transport, thus reducing toxic byproducts and improving cell growth under higher TAN concentrations. This finding is in good agreement with a previous study, in which the inactivation of antibiotic-specific porin decreases antibiotic uptake and susceptibility (40).

### New understanding regarding the mechanism of nitrification in methanotrophs

In previous works, Zahn et al. reported that CytL purified from *Methylococcus capsulatus* Bath converted NH_2_OH only to NO_2_^-^ (32); Versantvoorta et al. demonstrated that an HAO purified from *Verrucomicrobial* methanotroph *Methylacidiphilum fumariolicum* SolV could transform NH_2_OH to NO (27). Our study expands upon these insights by providing genetic evidence that the genes of *haoA*, *cytL* and *hcp* are involved in ammonia detoxification. Furthermore, biochemical analyses further revealed that CytL from *M. buryatense* 5GB1C could converts NH_2_OH to both N_2_O and NO_2_^-^ under aerobic condition. We also expressed Bath-CytL in *E. coli* and the purified Bath-CytL-6His was able to converted NH_2_OH to N_2_O and NO_2_^-^ under aerobic condition (data not shown). These results suggested that CytL from methanotrophs shares similar function with their counterparts in ammonia-oxidizing bacteria(23). Thirdly, the role of *hcp* in ammonia oxidation is worthy of further study. It has been suggested that Hcp can reduce NO to N_2_O in certain bacteria under anaerobic condition (43), while our results indicated that *hcp* was involved in ammonia toxicity under aerobic condition (FIG. 4E). In addition, overexpressing HAO increased the ammonium susceptibility (FIG. 4F), which probably was attributed to the toxicity of elevated NO. According to the catalytic mechanism observed in ammonia-oxidizing bacteria (22), CytL can convert NO to N_2_O in the presence of NH_2_OH. Consequently, the enhanced resistance to NH_4_Cl observed upon *cytL* overexpression supports this mechanism (FIG. 4G).

### Prospective work

If conversion of ammonia to NH_2_OH by pMMO is inevitable, it is important to note that the reduction of NH_2_OH and NO to N_2_O could lead to nitrogen loss and environment pollution. Therefore, a more effective strategy will focus on transforming NH_2_OH to NO and then to NO_2_^-^ in future. Encouragingly *M. buryatense* 5GB1C is capable of thriving on using NO_2_^-^ as nitrogen source (FIG. S1C).

## MATERIALS AND METHODS

### Bacterial Strains and Growth Conditions

The strains used in this study are listed in Table S1. The genome sequence of *M. buryatense* 5GB1C is available in GenBank with the accession number CP035467. *M. buryatense* 5GB1C was cultivated in NMS2 medium or AMS2 medium (36) under atmosphere of 25% methane in air. Liquid cultures were grown in sealed glass serum bottles, while plates were incubated in sealed jars at 30°C.

### Primers, Plasmids and Genetic Techniques

The primers and plasmids used in this study are listed in Table S2. Gene knockouts were carried out according to the method established by Liu et al (37). All enzymes used for DNA manipulation were purchased from Vazyme Biotech Co., Ltd (Nanjing, China) and used according to the manufacturer’s instructions. DNA sequencing was carried out by Sangon Biotech Co., Ltd (Shanghai, China).

### Measurement of Ammonia Oxidation Products

*M. buryatense* 5GB1C and its related mutants were first cultured in nitrate medium to the exponential phase. After centrifugation to collect the cells, the pellet was resuspended in 5 mL of medium containing NH_4_Cl to an OD_600_ of 0.8. The cells were then incubated 20 hours at 30°C with shaking at 180 rpm. NH_2_OH and NO_2_^-^ in the cell free supernatant were detected according to previous methods (26). N_2_O in the headspace was measured by a gas chromatography (GC7890B, Agilent, USA), equipped with flame ionization (FID) and electron capture (ECD) detectors. The ECD carrier gas consisted of high-purity N_2_ and argon-methane (Ar_2_:CH_4_= 95:5, v/v), with a detector temperature of 300°C and a column temperature of 50°C.

### Construction and Screening of a Transposon Mutagenesis Library

Transposon mutagenesis library of *M. buryatense* 5GB1C was constructed via triparental conjugation (36), with a donor strain of *E. coli* DH5α/pSC123 and a helper strain of *E. coli* DH5α/pRK600 (48, 49). The library was selected on NMS2 agar plates containing kanamycin (50 mg/L). Each colony was subsequently transferred to NMS2 and NMS2+NH_4_Cl plates and incubated for 3 days. Transposon insertion sites were identified using arbitrary PCR with primers Arb1, Arb2, and Arb3 (Table S2) (50).

### Analysis of TAN Adsorption

To assess the TAN adsorption capacity of wild-type strain and mutant strain Δ*porA*, cells first were grown to exponential phase in NMS2 medium, and then resuspended in AMS2 medium containing 300 µM NH_4_Cl to OD_600_ of 0.8. Cell free supernatants were collected at time points of 1, 3, 5, 10, and 20 minutes for quantification of TAN (51).

### Expression, Purification and Characterization of CytL

The gene encoding the protease DegP was deleted to generate strain *E. coli* BL21(DE3) Δ*degP*. CytL-6His was expressed using pET system and OmpA signal peptide in *E. coli* BL21(DE3) Δ*degP* (31). Meanwhile, the cluster of *ccmABCDEFGH* that is required for cytochrome *c* maturation was co-expressed in the same host (52). Expression of CytL-6His was induced with 0.1 mM IPTG at 16°C for 14 hours. CytL-6His was purified using Co-NTA affinity chromatography. The hydroxylamine-oxidizing activity of CytL-6His was measured in 8-mL septum-sealed cuvettes that contained 1 mL of 10.0 mM Tris-HCl buffer (pH 8.0), 10 µM CytL-6His, 1.0 mM NH_2_OH·HCl and 5 µM phenazine methosulfate (PMS). The reaction was incubated at 30°C.

### RT-qPCR

The cells of *M. buryatense* 5GB1C were cultured with KNO_3_ or NH_4_Cl as nitrogen source, then harvested at the early (OD_600_=0.4) and late (OD_600_=0.8) exponential phases. Total RNA was extracted using an RNA isolation kit (Sangon, No. B518655). Reverse transcription and removal of genomic DNA was performed with a HiScript III RT SuperMix (with gDNA wiper) (Vazyme, No. R323). cDNA was diluted and used as the template for qPCR, which was conducted in an Applied Biosystems 7500 real-time PCR system (Applied Biosystems, CA, USA) using ChamQ Universal SYBR qPCR Master Mix (Vazyme, No. Q711). Gene expression levels were normalized to 16S rRNA as the internal control, and relative expression was calculated using the 2⁻^ΔΔCT^ method.

### Fed-Batch Fermentation

Seed cultures were grown in three stages. Seed I in NMS2 medium and Seed II in AMS2 medium (2.0 mM NH_4_Cl) were both performed in 250-mL sealed glass serum bottles. Seed III in AMS2 medium (2.0 mM NH_4_Cl) was carried out in spinner bottles, and the composition and flow rate of mixed gas were the same as previously described (7). Fed-batch fermentation was conducted in 3-L stirred tank fermenters (Baoxing Bio-Engineering Equipment Co., Ltd, Shanghai, China) with a working volume of 2 L. Gas flow rate and temperature were maintained as described previously (7). pH was controlled around 8.0 with 1 mol/L H_2_SO_4_ and 2 mol/L NaOH. Dissolved oxygen was kept between 10-30% saturation by adjusting agitation speed between 200-700 rpm (FIG. S3A, B). Agitation speed was increased by 25-75 rpm each time, after 21.5 hours, the agitation speed was maintained at 550 rpm till the end. NH_4_Cl stock (214 g/L) was fed via peristaltic pump. The phosphate and trace element stock solution were intermittently fed at several time points (Table S4). TAN concentrations were measured using the Ammonia TNT plus Vial Test Kit (HACH, No. TNT-831). The specific growth rate (μ) was calculated from the slope of the natural log of OD_600_ versus time (35). Dry cell weight, methane uptake rate and total methane utilization were determined as previously described (7).

### Statistical analysis

Data analysis was performed using Student’s t-test or two-way ANOVA with GraphPad Prism software (version 8.3.0.538). *ns*, no significant difference; \**P* < 0.05; \*\**P* < 0.01; \*\*\**P* < 0.001; \*\*\*\**P* < 0.0001.

## ACKNOWLEDGMENTS

We thank Lisa Y. Stein for offering kind communication about the ammonia oxidation in methanotrophs. We thank J. Jiang, J Liu and L. Feng for their assistance in N_2_O detection. This work was supported by the National Natural Science Foundation of China (32270127, 32470117, 22178281) and the Science Fund for Distinguished Young Scholars of Shaanxi Province (2022JC-09).

## SUPPLEMENTAL MATERIAL

Figures S1 to S3; Tables S1 and S2.

## REFERENCES

1. Tveit A T, Hestnes A G, Robinson S L, Schintlmeister A, Dedysh S N, Jehmlich N, von Bergen M, Herbold C, Wagner M, Richter A. 2019. Widespread soil bacterium that oxidizes atmospheric methane. Proc Natl Acad Sci U S A 116: 8515–8524. 10.1073/pnas.1817812116

2. He L, Groom J D, Wilson E H, Fernandez J, Konopka M C, Beck D A, Lidstrom M E. 2023. A methanotrophic bacterium to enable methane removal for climate mitigation. Proc Natl Acad Sci U S A 120: e2310046120. 10.1073/pnas.2310046120

3. Conrado R J, Gonzalez R. 2014. Envisioning the bioconversion of methane to liquid fuels. Science 343: 621–623. doi:10.1126/science.1246929

4. Cantera S, Bordel S, Lebrero R, Gancedo J M, García-Encina P A, Muñoz R. 2019. Bio-conversion of methane into high profit margin compounds: an innovative, environmentally friendly and cost-effective platform for methane abatement. World. J. Microb Biot 35: 16. doi:10.1007/s11274-018-2587-4

5. Nguyen A D, Lee E Y. 2021. Engineered methanotrophy: A sustainable solution for methane-based industrial biomanufacturing. Trends Microbiol 39: 381–396. 10.1016/j.tibtech.2020.07.007

6. Abbadi S H E, Sherwin E, Brandt A, Luby S, Criddle C. 2022. Displacing fishmeal with protein derived from stranded methane. Nat Sustain 5: 47–56. 10.1038/s41893-021-00796-2

7. Gao Z, Guo S, Chen Y, Chen H, Fu R, Song Q, Li S, Lou W, Fan D, Li Y, Yang S, Gonzalez R, Fei Q. 2024. A novel nutritional induction strategy flexibly switching the biosynthesis of food-like products from methane by a methanotrophic bacterium. Green Chem 26: 7048–7058. 10.1039/d3gc04674e

8. Guo S, Song Q, Song X, Zhang C, Fei Q. 2024. Sustainable production of C50 carotenoid bacterioruberin from methane using soil-enriched microbial consortia. Bioresour Technol 412: 131415. 10.1016/j.biortech.2024.131415

9. Nguyen D T N, Lee O K, Lim C, Lee J, Na J-G, Lee E Y. 2020. Metabolic engineering of type II methanotroph, *Methylosinus trichosporium* OB3b, for production of 3-hydroxypropionic acid from methane via a malonyl-CoA reductase-dependent pathway. Metab Eng 59: 142–150. 10.1016/j.ymben.2020.02.002

10. Fu Y, Li Y, Lidstrom M E. 2017. The oxidative TCA cycle operates during methanotrophic growth of the Type I methanotroph *Methylomicrobium buryatense* 5GB1. Metab Eng 42: 43–51. 10.1016/j.ymben.2017.05.003

11. Risso C, Choudhary S, Johannessen A, Silverman J (2018) Methanotrophy Goes Commercial: Challenges, Opportunities, and Brief History. Methane Biocatalysis: Paving the Way to Sustainability, eds Kalyuzhnaya MG & Xing X (Springer, Cham), pp 293–298. 10.1007/978-3-319-74866-5_18

12. IMARC Group. Ammonium chloride pricing report 2024. https://www.imarcgroup.com/ammonium-chloride-pricing-report (Accessed 4 September 2024).

13. IMARC Group. Potassium nitrate pricing report 2024. https://www.imarcgroup.com/potassium-nitrate-pricing-report. (Accessed 4 September 2024).

14. Garg S, Clomburg J M, Gonzalez R. 2018. A modular approach for high-flux lactic acid production from methane in an industrial medium using engineered *Methylomicrobium buryatense* 5GB1. J Ind Microbiol Biotechnol 45: 379–391. 10.1007/s10295-018-2035-3

15. Meruvu H, Wu H, Jiao Z, Wang L, Fei Q. 2020. From nature to nurture: Essence and methods to isolate robust methanotrophic bacteria. Synth Syst Biotechnol 5: 173–178. 10.1016/j.synbio.2020.06.007

16. Fei Q, Liang B, Tao L, Tan E C, Gonzalez R, Henard C A, Guarnieri M T. 2020. Biological valorization of natural gas for the production of lactic acid: Techno-economic analysis and life cycle assessment. Biochem Eng J 158: 107500. 10.1016/j.bej.2020.107500

17. Stein L Y (2018) Proteobacterial Methanotrophs, Methylotrophs, and Nitrogen. Methane Biocatalysis: Paving the Way to Sustainability, eds Kalyuzhnaya MG & Xing X (Springer, Cham), pp 57–66. 10.1007/978-3-319-74866-5

18. Campbell M A, Nyerges G, Kozlowski J A, Poret-Peterson A T, Stein L Y, Klotz M G. 2011. Model of the molecular basis for hydroxylamine oxidation and nitrous oxide production in methanotrophic bacteria. FEMS Microbiol Lett 322: 82–89. 10.1111/j.1574-6968.2011.02340.x

19. Stein L Y, Klotz M G. 2011. Nitrifying and denitrifying pathways of methanotrophic bacteria. Biochem Soc Trans 39: 1826–1831. 10.1042/BST20110712

20. Kuypers M M, Marchant H K, Kartal B. 2018. The microbial nitrogen-cycling network. Nat Rev Microbiol 16: 263–276. 10.1038/nrmicro.2018.9

21. Stein L Y (2011) Heterotrophic Nitrification and Nitrifier Denitrification. Nitrification, eds Ward BB, Arp DJ, & Klotz MG (ASM Press, Washington), pp 95–114. 10.1128/9781555817145.ch5

22. Caranto J D, Lancaster K M. 2017. Nitric oxide is an obligate bacterial nitrification intermediate produced by hydroxylamine oxidoreductase. Proc Natl Acad Sci U S A 114: 8217–8222. www.pnas.org/cgi/doi/10.1073/pnas.1812827115

23. Caranto J D, Vilbert A C, Lancaster K M. 2016. *Nitrosomonas europaea* cytochrome P460 is a direct link between nitrification and nitrous oxide emission. Proc Natl Acad Sci U S A 113: 14704–14709. 10.1073/pnas.1611051113

24. Kozlowski J A, Price J, Stein L Y. 2014. Revision of N_2_O-producing pathways in the ammonia-oxidizing bacterium *Nitrosomonas europaea* ATCC 19718. Appl Environ Microbiol 80: 4930–4935. 10.1128/AEM.01061-14

25. Bédard C, Knowles R. 1989. Physiology, biochemistry, and specific inhibitors of CH_4_, NH_4_^+^, and CO oxidation by methanotrophs and nitrifiers. Microbiol Rev 53: 68–84. 10.1128/mr.53.1.68-84.1989

26. O’neill J G, Wilkinson J F. 1977. Oxidation of ammonia by methane-oxidizing bacteria and the effects of ammonia on methane oxidation. Microbiology 100: 407–412. 10.1099/00221287-100-2-407

27. Versantvoort W, Pol A, Jetten M S, van Niftrik L, Reimann J, Kartal B, Op den Camp H J. 2020. Multiheme hydroxylamine oxidoreductases produce NO during ammonia oxidation in methanotrophs. Proc Natl Acad Sci U S A 117: 24459–24463. 10.1073/pnas.201129911

28. Poret-Peterson A T, Graham J E, Gulledge J, Klotz M G. 2008. Transcription of nitrification genes by the methane-oxidizing bacterium, *Methylococcus capsulatus* strain Bath. ISME J 2: 1213–1220. 10.1038/ismej.2008.71

29. Sugden S, Lazic M, Sauvageau D, Stein L Y. 2021. Transcriptomic and metabolomic responses to carbon and nitrogen sources in *Methylomicrobium album* BG8. Appl Environ Microbiol 87: e00385–00321. 10.1128/AEM.00385-21

30. Guo K, Hakobyan A, Glatter T, Paczia N, Liesack W, Gilbert J A. 2022. *Methylocystis* sp. Strain SC2 Acclimatizes to Increasing NH_4_^+^ Levels by a Precise Rebalancing of Enzymes and Osmolyte Composition. mSystems 7: e00403–22 10.1128/msystems.00403-22

31. Adams H R, Krewson C, Vardanega J E, Fujii S, Moreno T, Sambongi Y, Svistunenko D, Paps J, Andrew C R, Hough M A. 2019. One fold, two functions: cytochrome P460 and cytochrome c′-β from the methanotroph *Methylococcus capsulatus* (Bath). Chem Sci 10: 3031–3041. 10.1039/c8sc05210g

32. Zahn J A, Duncan C, DiSpirito A A. 1994. Oxidation of hydroxylamine by cytochrome P-460 of the obligate methylotroph *Methylococcus capsulatus* Bath. J Bacteriol 176: 5879–5887. 10.1128/jb.176.19.5879-5887.1994

33. Bergmann D, Zahn J, Hooper A, Dispirito A. 1999. Cytochrome P460 Genes from the Methanotroph *Methylococcus capsulatus* Bath. J Bacteriol 180: 6440–6445. 10.1128/JB.180.24.6440-6445.1998

34. Kaluzhnaya M, Khmelenina V, Eshinimaev B, Suzina N, Nikitin D, Solonin A, Lin J-L, McDonald I, Murrell C, Trotsenko Y. 2001. Taxonomic characterization of new alkaliphilic and alkalitolerant methanotrophs from soda lakes of the Southeastern Transbaikal region and description of *Methylomicrobium buryatense* sp. nov. Syst Appl Microbiol 24: 166–176. 10.1078/0723-2020-00028

35. Gilman A, Laurens L M, Puri A W, Chu F, Pienkos P T, Lidstrom M E. 2015. Bioreactor performance parameters for an industrially-promising methanotroph *Methylomicrobium buryatense* 5GB1. Microb Cell Factories 14: 1–8. 10.1186/s12934-015-0372-8

36. Puri A W, Owen S, Chu F, Chavkin T, Beck D A, Kalyuzhnaya M G, Lidstrom M E. 2015. Genetic tools for the industrially promising methanotroph *Methylomicrobium buryatense*. Appl Environ Microbiol 81: 1775–1781. 10.1128/AEM.03795-14

37. Liu Y, He X, Zhu P, Cheng M, Hong Q, Yan X. 2020. *phes^AG^* based rapid and efficient markerless mutagenesis in *Methylotuvimicrobium*. Front Microbiol 11: 441. 10.3389/fmicb.2020.00441

38. de la Torre A, Metivier A, Chu F, Laurens L M, Beck D A, Pienkos P T, Lidstrom M E, Kalyuzhnaya M G. 2015. Genome-scale metabolic reconstructions and theoretical investigation of methane conversion in *Methylomicrobium buryatense* strain 5G (B1). Microb Cell Factories 14: 1–15. 10.1186/s12934-015-0377-3

39. Yan X, Chu F, Puri A W, Fu Y, Lidstrom M E. 2016. Electroporation-based genetic manipulation in type I methanotrophs. Appl Environ Microbiol 82: 2062–2069. 10.1128/AEM.03724-15

40. Vergalli J, Bodrenko I V, Masi M, Moynié L, Acosta-Gutiérrez S, Naismith J H, Davin-Regli A, Ceccarelli M, van den Berg B, Winterhalter M, Pagès J-M. 2020. Porins and small-molecule translocation across the outer membrane of Gram-negative bacteria. Nat Rev Microbiol18: 164–176. 10.1038/s41579-019-0294-2

41. Dam S, Pagès J-M, Masi M. 2018. Stress responses, outer membrane permeability control and antimicrobial resistance in enterobacteriaceae. Microbiology 164. 10.1099/mic.0.000613

42. Wolfe M T, Heo J, Garavelli J S, Ludden P W. 2002. Hydroxylamine reductase activity of the hybrid cluster protein from *Escherichia coli*. J Bacteriol 184: 5898–5902. 10.1128/jb.184.21.5898-5902.2002

43. Wang J, Vine C E, Balasiny B K, Rizk J, Bradley C L, Tinajero-Trejo M, Poole R K, Bergaust L L, Bakken L R, Cole J A. 2016. The roles of the hybrid cluster protein, Hcp and its reductase, Hcr, in high affinity nitric oxide reduction that protects anaerobic cultures of *Escherichia coli* against nitrosative stress. Mol Microbiol 100: 877–892. 10.1111/mmi.13356

44. Hagen W. 2022. Structure and function of the hybrid cluster protein. Coord Chem Rev 457: 214405. 10.1016/j.ccr.2021.214405

45. Stein L Y, Lidstrom M E. 2024. Greenhouse gas mitigation requires caution. Science 384: 1068–1069. 10.1126/science.adi0503

46. Molins-Legua C, Meseguer-Lloret S, Moliner-Martinez Y, Campíns-Falcó P. 2006. A guide for selecting the most appropriate method for ammonium determination in water analysis. Trends Analyt Chem 25: 282–290. 10.1016/j.trac.2005.12.002

47. Emerson K, Russo R C, Lund R E, Thurston R V. 1975. Aqueous ammonia equilibrium calculations: Effect of pH and temperature. J Fish Res Board Can 32: 2379–2383. 10.1139/f75-274

48. Simon R, Priefer U, Pühler A. 1983. A broad host range mobilization system for in vivo genetic engineering: Transposon mutagenesis in gram negative bacteria. Nat Biotechnol 1: 784–791. 10.1038/nbt1183-784

49. Kessler B, de Lorenzo V, Timmis K N. 1992. A general system to integrate *lacZ* fusions into the chromosomes of gram-negative eubacteria: regulation of the *Pm* promoter of the *TOL* plasmid studied with all controlling elements in monocopy. Mol Gen Genet 233: 293–301. 10.1007/BF00587591

50. Judson N, Mekalanos J J. 2000. TnAraOut, A transposon-based approach to identify and characterize essential bacterial genes. Nat Biotechnol 18: 740–745. 10.1038/77305

51. Bower C E, Holm-Hansen T. 1980. A salicylate–hypochlorite method for determining ammonia in seawater. Can J Fish Aquat Sci 37: 794–798. 10.1139/f80-106

52. Arslan E, Schulz H, Zufferey R, Künzler P, Thöny-Meyer L. 1998. Over production of the Bradyrhizobium japonicum c-type cytochrome subunits of the cbb_3_ oxidase in *Escherichia coli*. Biochem Biophys Res Commun 251: 744–747. 10.1006/bbrc.1998.9549

